# Insights into Digit Evolution from a Fate Map Study of the Forearm

**DOI:** 10.1101/2023.08.29.555165

**Authors:** JDH Oh, DDZ Saunders, L McTeir, M Jackson, JD Glover, JJ Schoenebeck, LA Lettice, MG Davey

## Abstract

The cellular and genetic networks which contribute to the development of the zeugopod, (radius and ulna of the forearm, tibia and fibula of the leg) are not well understood, although these bones are susceptible to loss in congenital human syndromes and to the action of teratogens such as thalidomide. Using a new fate mapping approach in transgenic chickens, we show that there is a small contribution of *SHH* expressing cells to the posterior ulna, posterior carpals and digit 3. We establish that while the majority of the ulna develops in response to paracrine SHH signaling in both the chicken and mouse, there are differences in the contribution of *SHH* expressing cells to other tissues of the zeugopod between these two species as well as between the chicken ulna and fibula. This is evidence that although zeugopod bones are clearly homologous according to the fossil record, the zeugopod bones of the wing and leg are formed by subtly different signalling and patterning events during embryonic development, which can be used to understand the shaping of the bird wing skeleton during the evolution of powered flight.

## Introduction

Limbs first form as small paired forelimb or hindlimb buds growing from the flank of a developing embryo (Tickle, 2015). The mesodermal cell component of the limb bud, derived from the lateral plate mesoderm (Gros and Tabin, 2014), forms the majority of the limb skeleton from the proximal shoulder/pelvic girdle to the digit tips. The cells that make up the early limb look homogenous but fate maps of the early chicken wing bud show that at stage 20HH (Hamburger and Hamilton, 1951), mesodermal-derived cells within specific areas are already fated to form either the shoulder/pelvic girdle, stylopod (humerus/femur) or zeugopod (radius and ulna, tibia/fibula), and by HH24, the autopod (digits; Dudley et al., 2002; Nomura et al., 2014; Sato et al., 2007; Saunders, 1948; Vargesson et al., 1997). Within the autopod, the origin, number and signalling pathways which pattern the antero-posterior identity of digits have been well studied (Harfe et al., 2004; Tamura et al., 2011; Towers et al., 2008; Towers et al., 2011; Zhu et al., 2022). Although the specification of the zeugopod region within the proximo-distal axis of limb bud has been examined (Dudley et al., 2002; McCusker and Rosello-Diez, 2022; Rosello-Diez et al., 2011; Sato et al., 2007) how two bones with different antero-posterior identities, the anterior radius and posterior ulna, develop from this area has not been thoroughly investigated. Human conditions highlight the separate identities of these bones in that there are notable differences between conditions where either the radius or ulna is lost. Radial deficiency is more common than ulnar deficiency, even in thalidomide cases where it is more common to observe the loss of entire proximo-distal segments. Unlike radial deficiency, ulnar deficiency is rarely associated with systemic syndromes (Bednar et al., 2009).

The zeugopod is, however, subject to many of the same patterning mechanisms as the autopod and parallels between these parts of the limb can be drawn, specifically between the antero-posterior axis patterning by SHH and FGF pathways (Chiang et al., 2001; Mariani et al., 2008). A loss of FGF signaling in the mouse or inhibition of cell proliferation in the chicken limb bud causes a loss of anterior digits and the radius, evidence that these tissues are dependent on cell proliferation driven by FGF signaling (Mariani et al., 2008; Towers et al., 2008; Towers and Tickle, 2009). In the chicken wing, *SHH* is expressed in the mesoderm-derived organiser of the limb, the ‘Zone of Polarising Activity’ (ZPA), from stage 18HH and it is thought that the relative balance of paracrine and autocrine SHH signaling, along with cell proliferation, is central to establishing both digit number and identity (Towers et al., 2008; Zhu et al., 2022). In the human, mouse or chicken, a loss of SHH causes a loss of posterior digits and a loss of the ulna (Chiang et al., 2001; Ianakiev et al., 2001; Ros et al., 2003; Towers et al., 2008), demonstrating that SHH signaling is required for posterior limb identity in either the zeugopod or autopod and that the ulna is a SHH-dependent bone. In addition there is a distinction between the derivatives of cells expressing *SHH* within the ZPA organiser and subject to autocrine SHH signalling and those which are patterned by the ZPA organiser, receiving paracrine SHH signals. In the mouse, *Shh* expressing cells from the ZPA contribute to digits 3-5 as well as the ulna (Harfe et al., 2004; Scherz et al., 2007), indicating that a portion of the ulna is patterned by autocrine Shh signaling as well as paracrine signaling (Ahn and Joyner, 2004). The contribution of *SHH* expressing cells to the ulna has not been examined in the chick, although unlike the 5-fingered mouse, *SHH* expressing cells do not contribute to any of the three digits of the wing (Towers et al., 2011). This has been used as evidence to determine which two digits birds lost during evolution towards powered flight (Tamura et al., 2011; Towers et al., 2011; Xu and Mackem, 2013), an important paradigm in the study of evolutionary development (Evo-Devo).

The evolution of the bird wing, in particular understanding which two digits were ‘lost’ and which three remain in the modern tridactyly wing, is studied both to understand the context of the bird wing as a model of vertebrate limb development and morphological evolution (Brusatte, 2017; Richardson et al., 2009). The focus on the majority of the research in this area has been to understand which of the three bird digits are homologous to a five of a pentadactyl limb, such as a mouse, human or basal archosaur, an example of which is the basal tetrapod *Westlothiana* (Smithson et al., 2011) from which all limbs’ pattern arose.

There are conflicting interpretations of digit homology due to an incomplete fossil record and confounded by an ambiguity in assigning a universal digit identity to either the three bird digits, using either adult or embryological data (Burke and Feduccia, 1997; Chatterjee, 1998; de Bakker et al., 2013; de Bakker et al., 2021; Hinchliffe and Hecht, 1984; Kawahata et al., 2019; Larsson and Wagner, 2002; Richardson, 2012; Salinas-Saavedra et al., 2014; Stewart et al., 2019; Tamura et al., 2011; Towers, 2018; Towers et al., 2008; Towers et al., 2011; Vargas and Fallon, 2005; Welten et al., 2005; Woltering and Duboule, 2010; Xu and Mackem, 2013; Xu et al., 2014). In these studies, evolutionary anatomical changes in the zeugopod bones, have been overlooked as homology of the radius and ulna is easily assigned and both are clearly present throughout the fossil record. Rather, the emphasis has been that morphology of the carpals and digits has evolved distal to the ‘unchanging’ bony anatomy of the forearm, the radius and ulna (Fig. 1D). This is embodied in the a foundation principal, the ‘primary limb axis’ hypothesis (Salinas-Saavedra et al., 2014; Shubin and Alberch, 1986), which emphasises the line of conserved morphology that includes the humerus and ulna around which distally digits have evolved. How palaeontological, anatomical and embryological data have been interpreted has led to the development of the ‘frame-shift’ and ‘axis-shift’ hypotheses (Xu and Mackem, 2013).

**Figure 1.**
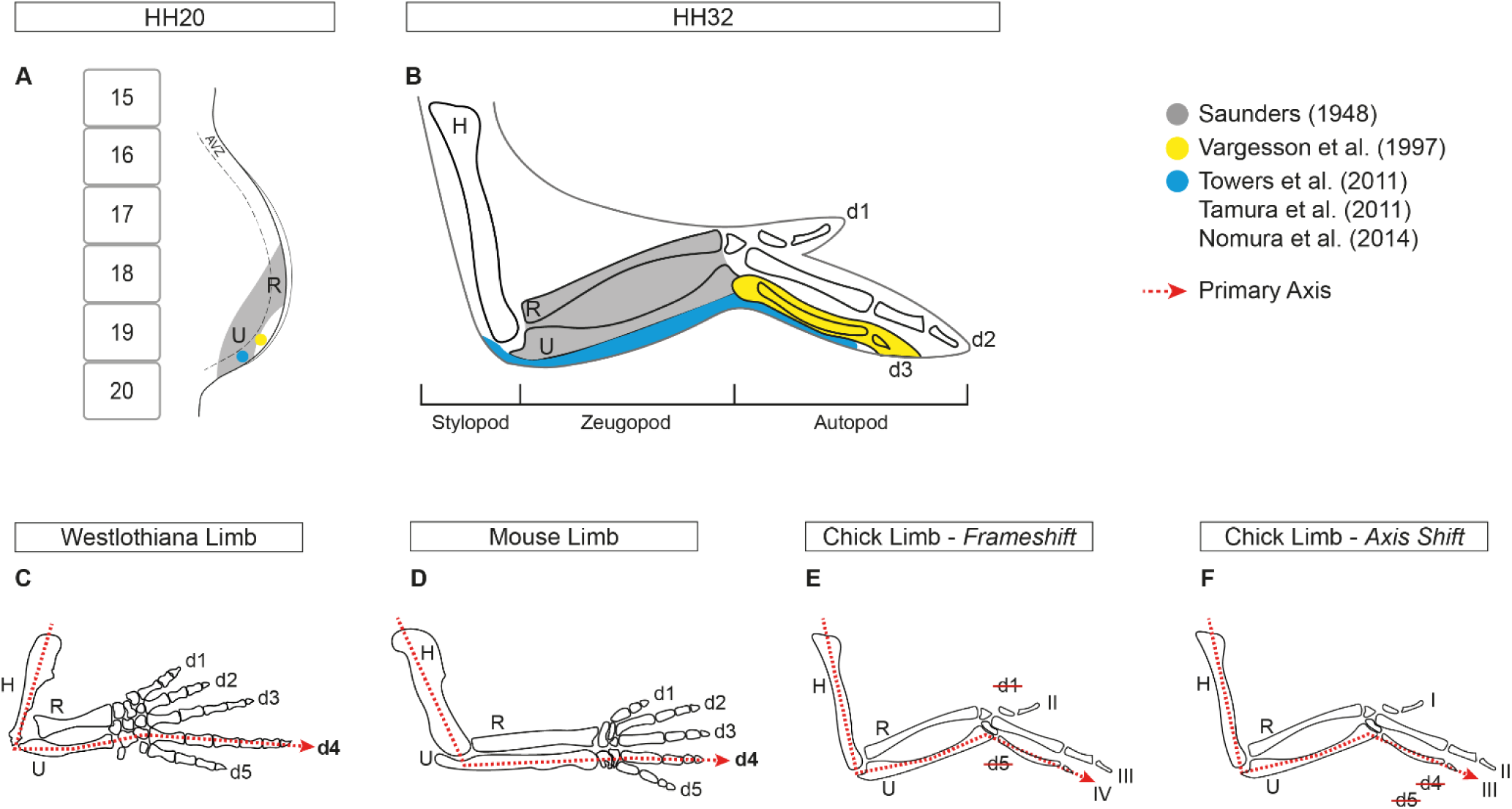
Summary of published fate maps and hypotheses for digit loss in chicken. Amalgamation of previous fate map studies of the chick wing with (**A**) showing the stage 20HH chick wing bud with somites and avascular zone as reference and (**B**) showing the stage 32HH chick wing. Grey shading derived from Saunders (1948) and yellow shading from Vargesson et al. (1997). Blue shading show an agreement of results from Towers et al. (2011), Tamura et al. (2011) and Nomura et al. (2014). Diagram of the primary axis represented by a red dotted line going through the humerus, ulna and digit 4 in (**C**) the Westlothiana limb and (**D**) mouse limb. (**E**) Schematic of the frameshift hypothesis, in which the primary axis continues to course through the ulna and digit 4 in the chick wing, with a loss of digits 1 and 5. (**F**) Schematic of the axis shift hypothesis, in which the primary axis has shifted and now courses through digit 3 in the chick wing. <u>Abbreviations</u> HH: Hamburger Hamilton. AVZ: Avascular zone. H: Humerus. R: Radius. U: Ulna. d: digit.

The ‘frame-shift’ model (Fig. 1E), primarily based on embryological evidence such as the development of *SOX9*+ digit primordia, proposes that the primary axis is maintained and the ulna-digit 4 articulation remains unchanged, but that a modified digit 4 takes on a morphological identity of a digit III through a homeotic transformation, thereby concluding that digit 1 and 5 are lost (de Bakker et al., 2013). Alternatively based on both fossil and embryological data, specifically the contribution of *SHH* expressing cells to the digits as a indicator of lineage, the ‘axis-shift’ model (Fig. 1F) suggests that the articulation between the primary axis/ulna shifts from digit 4 to digit 3, but does not account for how the change in this relationship might have occurred (Towers et al., 2011). A limitation of all these studies has been a lack of analysis of the bones proximal to the digits although analysis of the carpals suggests that these bones, articulating the zeugopod with the autopod, have been even more radically altered than the digits (Botelho et al., 2014). We propose that understanding developmental events which pattern the limb proximal to the digits, including the contribution of *SHH* expressing cells to elements of the posterior bird forelimb and carpals, is central to understanding the evolution of the avian primary limb axis and digits that articulate with it. We therefore sought to identify the exact location of the ulna anlage and explore its relation to the ZPA using a new anatomical approach to fate mapping in the developing chicken embryo. We show that, like the mouse, SHH ZPA cells contribute to the chicken ulna, carpals and digit 3 cartilage in a developmental stage dependent manner, demonstrating an embryological relationship between these skeletal elements.

## Results

### The ulna arises from a discrete area within the chick limb bud

To locate the area from which the ulna is specified in the stage 20HH chick wing bud, we employed a novel fate map technique that utilises the Chameleon cytbow chicken line in conjunction with TAT-Cre recombinase (Davey et al., 2018). Initially, all cells in the Chameleon chick embryo ubiquitously express nuclear H2B-eBFP2. Addition of beads soaked with TAT-Cre recombinase to the Chameleon chick embryo induces recombination at the cytbow transgene, deleting the nuclear H2B-eBFP2 and allowing expression of one of the three fluorescent proteins: eYFP, tdTomato or mCFP (accompanying Bio-Protocol paper; Saunders et al.). The action of TAT-Cre protein in the developing embryo is both highly localised to the area of application and transient, lasting less than a minute, resulting in small discrete induction of stable fluorescence expression which can be subsequently assessed clonally.

As the ulna is known to be dependent on SHH signaling, we first examined the fate of the presumptive zeugopod forming region at 20HH as identified by Saunders (1948; Fig. 1A, B). With Saunders’ map as a guide, beads soaked in TAT-Cre recombinase were inserted around the ulnar area of the presumptive zeugopod forming region in 20HH Chameleon chicken limb buds to determine that the ulna arises from cells in the distal limb bud, parallel to the anterior half of somite 19 (Fig. 2A, B). This region of the limb lies above the *SHH* expressing ZPA cells but expresses *PTCH1*, a hedgehog receptor whose expression is induced by the ligand, demonstrating the area is subject to paracrine SHH signaling (Fig. 2C, C’’). Stage 33HH wings were subsequently analysed for anatomical distribution of fluorescent cells, which were found to be located in the ulna, posterior carpals and digit 3 (n=3; Fig. 2D-J). Beads placed either more proximally or within the ZPA parallel to the posterior half of somite 19, which expresses both *SHH* and *PTCH1,* did not result in fluorescent labelling of the ulna (n=5; Supplementary Fig. 1).

**Figure 2.**
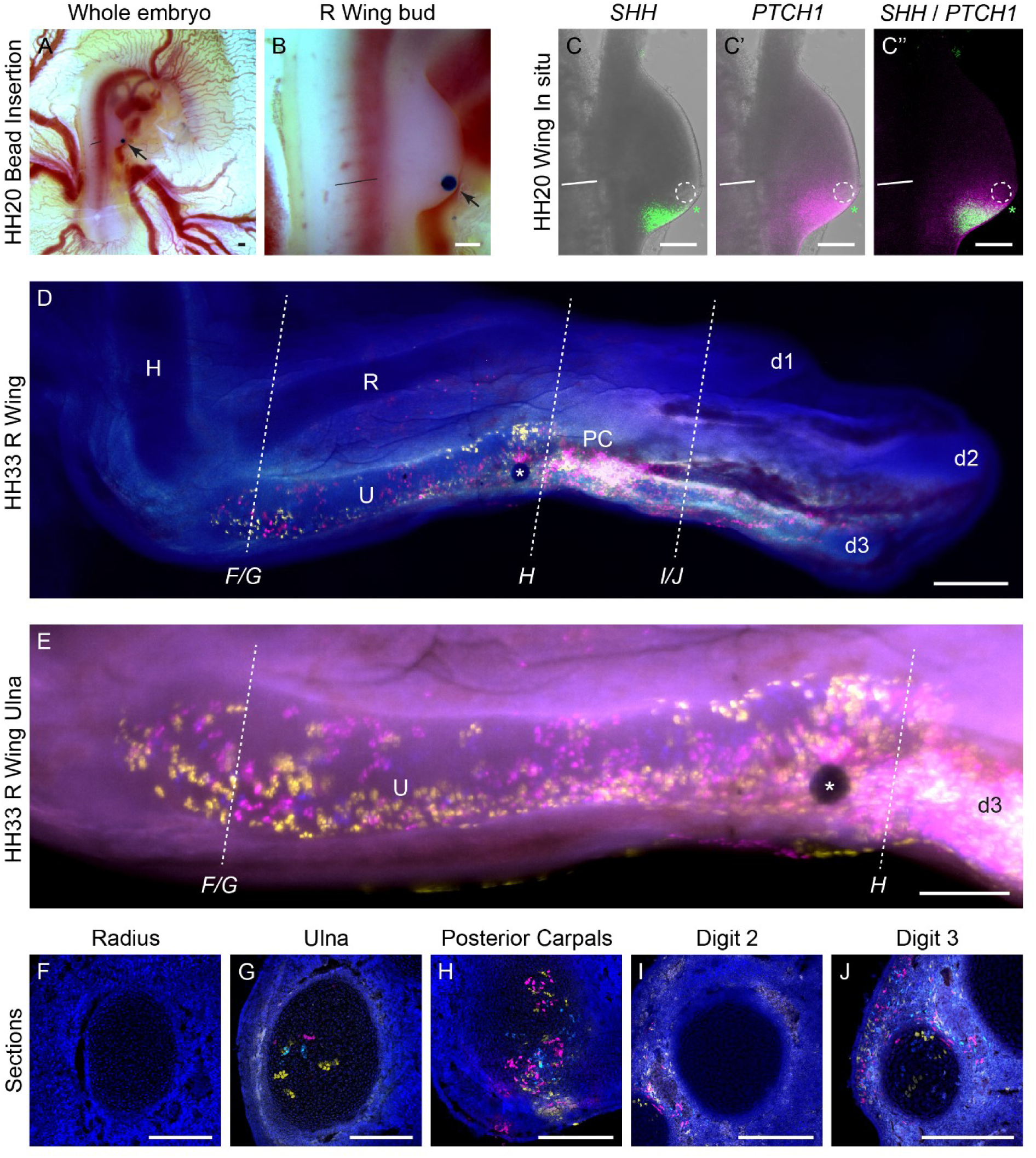
**Fate map of chick ulna using Chameleon chickens.** Placement of beads soaked in TAT-Cre (arrow) that maps the ulna in stage 20HH Chameleon chick wing buds shown in (**A**) whole embryo and (**B**) and at higher magnification. Stage 20HH chick wing buds with inert bead (dashed white circle) inserted in anterior half of somite 19, same as (**B**) then HCR in situ hybridisation performed with (**C**) *SHH* and (**C’**) *PTCH1* both shown against brightfield. Merge of *SHH* and *PTCH1* shown in (**C’’**). Asterisk denotes the anterior-most edge of the ZPA. Straight black and white lines denote the anterior-most edge of somite 19. Chameleon stage 33HH chick wing showing fluorescent cells (magenta, yellow and cyan with white indicating overlap) in the ulna and digit 3 (**D**) which were recombined on exposure to TAT-Cre delivered by bead as per (**B**). Close-up of the same limb with focus on ulna (**E**). Dashed lines in (**D**) and (**E**) denote where sections of the radius (**F**), ulna (**G**), posterior carpals (**H**), digit 2 (**I**) and digit 3 (**J**) were taken. White asterisk denotes location of bead. <u>Abbreviations</u> HH: Hamburger Hamilton. AVZ: Avascular zone. H: Humerus. R: Radius. U: Ulna. PC: Posterior carpals. d1/2/3: digit 1/2/3. All scale bars = 200μm

Closer analysis of limbs labelled at the anterior half somite 19 (*SHH*-/*PTCH1*+), demonstrated that fluorescent cells spanned the length of the ulna (Fig. 2D-G) and were largely contained within the ulna cartilage (Fig. 2E-G) indicating that cells within 50µm of the bead at stage 20HH contributed to the entire length of the ulna. Sections showed no labelled cells in the radius (Fig. 2F) and few cells in the ulnar perichondrium or adjacent soft tissues (n=3/3, Fig. 2E-G). In addition to the ulna, the cartilage of posterior carpals (Fig. 2H) and the cartilage of digit 3 (Fig. 2J) also contained fluorescent cells, as well as soft tissue adjacent to the cartilage of digit 2 (Fig. 2I; n=3/3).

### *SHH* expressing cells make a small contribution to the ulna in a stage-dependent manner

Our fate mapping approach, like others before, creates small and discrete clones of labelled cells. While excellent for generating fate maps with high spatial resolution, it does not demonstrate the fate of all the cells in a specific region. For example, no single bead application labelled all the cells of an ulna (Fig. 2D). Until recently it has been generally presumed that each bone forms from one area, or primordia, which can be sufficiently represented by small clones of labelled cell in fate napping approaches. Using a different fate mapping approach in mouse, it has recently been shown that rather than expanding over time and differentiating in a proximal to distal order from one primordia, limb bones, including the ulna, form piecemeal with different parts of the bone differentiating at different times but eventually forming one entity (Markman et al., 2023), although this evidence is not inconsistent with individual bones arising from one primordia. To assess if the area we had identified was able to generate all the cells of the ulna, we undertook homotopic grafting of the presumptive ulna primordia between stage 20/21HH eGFP and dtTomato transgenic chicken embryos. Distal wing mesenchyme grafts from dtTomato stage 20HH limbs, corresponding to the anterior of somite 19 and approximately 150µm by 150µm in size (Fig. 3A), were grafted into the equivalent area in eGFP embryos (Fig. 3B). To confirm that grafts were correctly taken from the *SHH*-/*PTCH1*+ domain we observed examined gene expression in donor limbs after grafts were excised, via HCR RNA in situ hybridisation and confirmed that all grafts originated from the *SHH*-/*PTCH1*+ presumptive ulna primordia (Fig. 3M-M’’’). qRT-PCR was used to assess expression in mock grafts from the presumptive ulna primordia which were also found to be *SHH*-/*PTCH1*+ (Fig. 3L). tdTom grafts of the presumptive ulna primordia gave rise to the cartilage of the ulna and carpals (n=7/7; Fig. 3C-D’) and digit 3 (n=5/7) in host eGFP embryos. Unlike labelling of the ulna primordia via TAT-Cre application, contribution to the entire length of the ulna was dependent on graft size as smaller grafts only gave rise to the distal ulna, carpals and digit (n=4/7). However, this demonstrates that the cells which generate the ulna at stage 20/21HH come from within the distal *SHH*-/*PTCH1*+ domain, outside of the *SHH* expressing ZPA.

**Figure 3.**
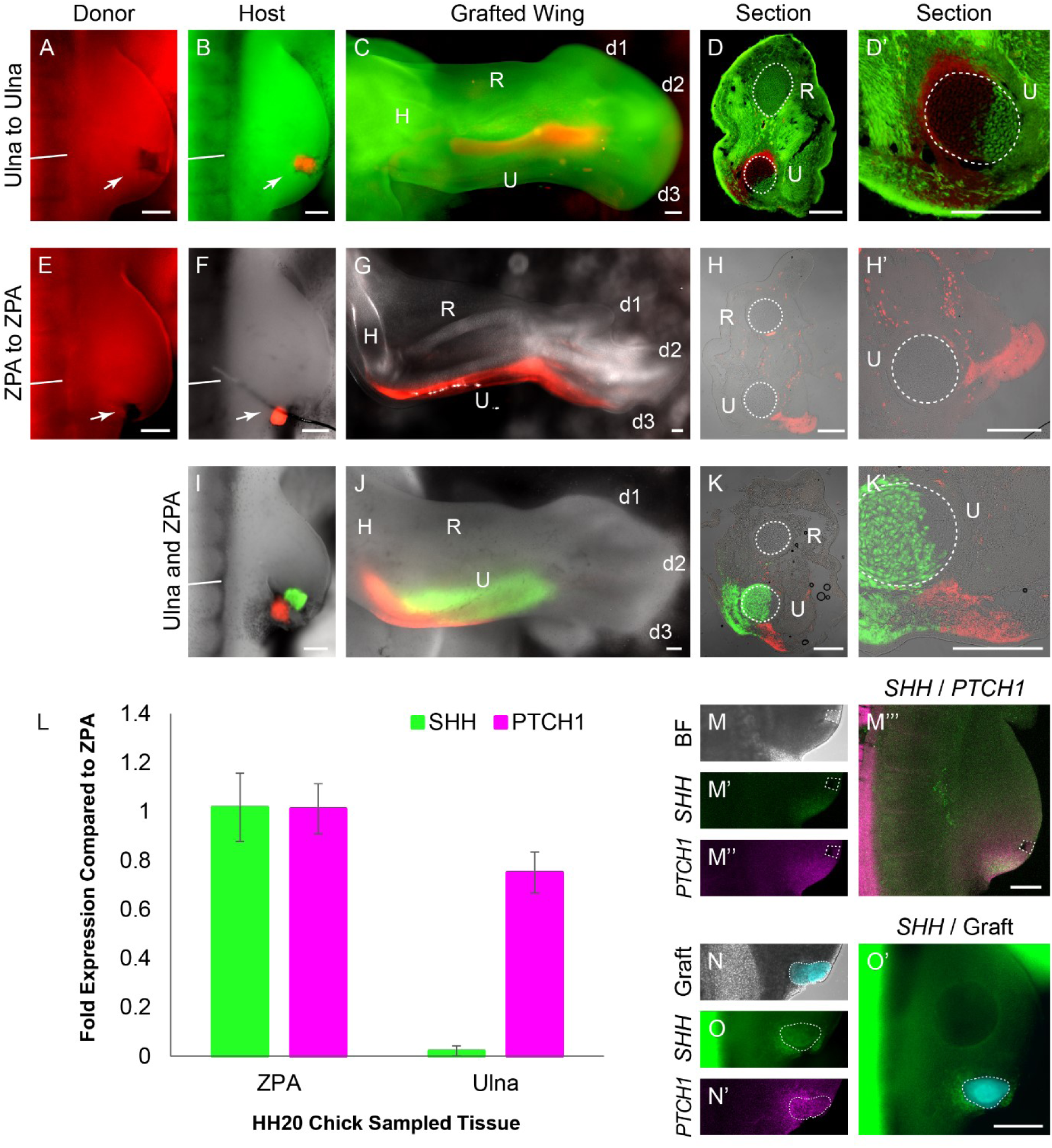
**ZPA lineage in chick wing in association with *SHH* and *PTCH1* expressions** Confirmation of Chameleon results by homotopic grafting of presumed ulna from stage 20HH tdTom chick wing bud (**A**) to eGFP chick wing bud (**B**). tdTom cells contribute to the entire length of the ulna as shown in (**C**) with sections (**D**, **D’**) confirming tdTom cells in cartilage. ZPA lineage determined through homotopic grafting of ZPA from stage 20HH tdTom chick wing bud (**E**) to wild-type chick wing bud (**F**). tdTom cells do not contribute to the ulna as shown in wholemount (**G**), confirmed with sections (**H**, **H’**). White arrows indicate graft donor and host site. Homotopic double grafts with ZPA derived from a tdTom chick wing bud and presumed ulna from an eGFP chick wing bud grafted into wild-type wing bud (**I**). Subsequent wholemount (**J**) and sections (**K**, **K’**) show only eGFP cells contribute to ulna. Dashed white circles outline the ulna and radius in sections. qRT-PCR for *SHH* (green) and *PTCH1* (magenta) performed for 20HH ZPA, ulna and radius primordia (**L**). Close-up of 20HH chick wing with either presumed ulna excised (dashed white box) (**M**) then HCR in situ hybridisation performed with *SHH* (**M’**) and *PTCH1* (**M’’**). The same limb with merge of *SHH* and *PTCH1* (**M’’’**). Close-up of 20HH chick wing with eGFP ZPA grafted into wild-type host (dashed white line) (**N**) then HCR in situ hybridisation performed with *SHH* (**O**) and *PTCH1* (**N’**) with merge of *SHH* and graft in (**O’**). (**N**, **N’**) and (**O**, **O’**) are of the same limb but imaged with confocal and fluorescent zoom microscopes, respectively. Straight white lines denote the anterior-most edge of somite 19. <u>Abbreviations</u> H: Humerus. R: Radius. U: Ulna. d1/2/3: digit 1/2/3. BF: Brightfield. All scale bars = 200μm.

It had previously been reported that in mice, the ulna arises from *SHH* expressing cells (Harfe et al., 2004). In 20/21HH homochronic *SHH*-/*PTCH1*+ domain grafts, complete ulna labelling was observed in 3/7 samples. To confirm that stage 20/21HH chick ZPA cells (*SHH*+/*PTCH1*+) do not contribute to the ulna, we performed homotopic ZPA grafts from 20HH dtTom or eGFP embryos (Fig. 3E) to either eGFP or non-transgenic chick wings (Fig. 3F). RT-qPCR was used to confirm expression of *SHH* and *PTCH1* in ‘mock’ ZPA grafts, with around a 47 fold decrease in *SHH* of the ulna compared to the ZPA (p<0.05, Fig. 3L).

HCR RNA in situ hybridisation was used to assess expression of *SHH* and *PTCH1* in host embryos containing grafts, confirming that grafted tissue originating from the ZPA (*SHH*+/*PTCH1*+) was grafted into the ZPA region (*SHH*+/*PTCH1*+) in mock grafting experiments (n=4, Fig. N-O’). All ZPA-ZPA stage 20HH grafts (n=3) contributed to the posterior mesenchyme of the limb at stage 33HH but not the ulna, confirming that the stage 20HH ZPA does not contribute cells to the ulna (Fig. 3G-H’).

To explore the interaction between ZPA cells and the ulnar primordium, both the ZPA (tdTom) and the region giving rise to the ulna (eGFP) were transplanted together into wildtype stage 20HH limb buds (Fig. 3I). At stage 33, there was no mixing of eGFP and tdTomato cells in all limbs (Fig. 3J; n=3). Only eGFP cells (*SHH*-/*PTCH1*+) were within the ulna cartilage and ZPA derived tdTom cells remained strictly outside of the cartilage (Fig. 3K, Supplementary Fig. 2).

These results demonstrate that the ulnar primordium is spatially defined, consistent with the original chicken limb fate maps of Saunders (1948) and Summerbell (1974), but *SHH* expressing cells do not contribute to the ulna in the chicken, as previously described in mouse (Harfe et al., 2004).

To further explore potential differences in ulna specification between chicken and mouse we re-examined the SHH reporter mouse SHH^tm1(EGFP/cre)Cjt^ (Harfe et al., 2004; Scherz et al., 2007). In combination with data from online resources (Baldarelli et al., 2021; J:184579; Shh Embryo 3 E15.5; https://images.jax.org/webclient/img_detail/17489/) we confirm that, while SHH expressing cells do contribute to the ulna and posterior mesenchyme of the zeugopod, localisation is primarily in the distal ulna and is far less extensive than the contribution to digit 4 and 5 (Harfe et al., 2004; Fig. 4A-E).

**Figure 4.**
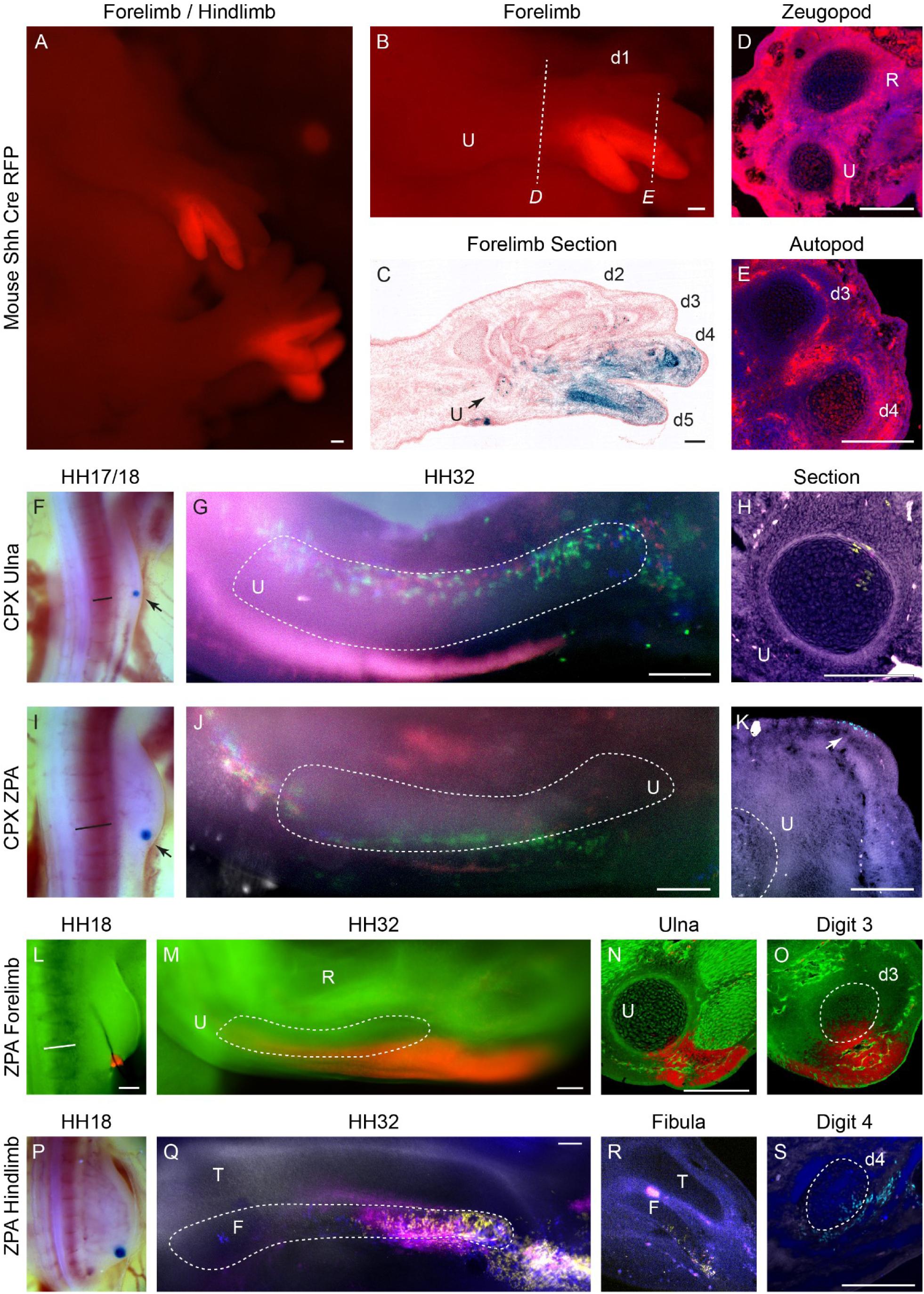
**ZPA lineage in forelimbs and hindlimbs of mice and HH18 chicken.** Right forelimb and hindlimb of an embryo carrying both SHH^tm1(EGFP/cre)Cjt^ and a cre inducible tdRFP reporter with RFP expression highlighting cells in the *SHH* lineage (**A**). Same mouse with close-up of forelimb (**B**) with RFP in distal ulna and digits 4 and 5. Longitudinal section from J:184579 (see main text) with blue showing *SHH* lineage (**C**). Black arrow points to the distal ulna. Dashed lines in (**B**) denote where sections of the zeugopod (**D**) and autopod (**E**) were taken with RFP in the cartilage of the ulna and digit 4. TAT-Cre bead placement (arrow) for the ulna in the Chameleon 17HH wing bud (**F**) and for the ZPA in the Chameleon 18HH wing bud (**I**) with subsequent wholemounts in (**G**) and (**J**). Sections show that fluorescent cells are within ulna cartilage for bead placed outside the ZPA (**H**) but also for beads placed within the ZPA (**K**). Homotopic tdTom ZPA to eGFP grafts in 18HH wing buds (**L**) to confirm contribution of tdTom to a minority of ulna cartilage but also to digit 3 cartilage shown in wholemount (**M**) and sections (**N**, **O**). TAT-Cre bead placement in the ZPA of Chameleon 18HH hindlimb (**P**) with wholemount (**Q**) and sections (**R**, **S**) showing ZPA lineage in the fibula and digit 4. Straight black and white lines denote the anterior-most edge of somite 19. Abbreviations HH: Hamburger Hamilton. R: Radius. U: Ulna. T: Tibia. F: Fibula. d1/2/3/4/5: digit 1/2/3/4/5. All scale bars = 200μm

Apical ectoderm ridge excision experiments of both Saunders (1948) and Summerbell (1974) suggest that the proximal chicken ulna is specified before stage 20HH, so we therefore sought to establish if the difference between mouse and chicken data could be resolved by undertaking ZPA grafts earlier in development. We implanted TAT-Cre beads into the distal limb mesenchyme of stage 18HH limbs at the axial level of anterior somite 19 (Fig. 4F) and more posteriorly into the ZPA (Fig. 4I). Localisation of fluorescent clones were substantially different between the experiments; beads placed in the “ulna region” at anterior somite 19 resulted in fluorescent labelling in the stylopod, the cartilage of the ulna and a small contribution to the autopod (n=5/5; Fig. 4G, H). Beads placed in the ZPA, however, labelled the stylopod, posterior limb mesenchyme of the zeugopod and autopod, but not the ulna cartilage (n=4; Fig.4J, K). Additionally, we undertook homotopic grafting of dtTom ZPA grafts to stage 18HH eGFP embryos (Fig. 4L). In this instance we did find that dtTom ZPA grafts made a small contribution to posterior ulna and digit 3 (n=3/3; Fig. 4M-O), two of which also contributed to the full length of the ulna. This work reconciles the origin of the ulna between mouse and chicken, showing both have a small contribution of *SHH* expressing cells along the posterior side of the ulna and digit 3 cartilages.

The chick hindlimb comprises of four digits and is considered to be a closer representative of the pentadactyl limb of mice. The fourth digit of the chick leg is predominantly descended from ZPA cells whilst the three anterior digits are not ZPA descendants, echoing ZPA contributions to digits in mice (Towers, 2018). This suggests that there has been a loss of the fifth digit in birds, which is reflective of fossil records of theropods. The zeugopod of the chick hindlimb, fibula and tibia, are analogous to the ulna and radius of the forelimb, respectively. To examine if the fibula, like the ulna of the mouse and chicken also arises predominantly from *SHH*-/*PTCH1*+ cells, we implanted Tat-Cre beads into the ZPA of 18HH hindlimb buds (Fig. 4P) and found surprisingly that the resulting fluorescent clones contributed to the distal two-thirds of the fibula, the fourth metacarpal and phalanges of digit 4 (Fig. 4Q-S). This suggests a much larger contribution of SHH expressing cells form the fibula than the ulna and demonstrates that even between the two posterior zeugopod bones of birds (i.e. ulna and fibula), there is a considerable difference in the cellular lineages which comprise them.

## Discussion

Fate mapping approaches have been fundamental in developmental biology and the chicken embryo has been particularly useful in developing anatomical and temporal fate maps of developing tissues due to its anatomical accessibility. Limitations in technology, however, have also limited insights that can be made. Here we demonstrate a new anatomical approach to fate-mapping, which utilises topically applied TAT-Cre to a transgenic chicken containing a Cre-inducible transgene. This approach faithfully recreates and improves on fate maps of the chicken limb made by Saunders (1948), Vargesson et al (1997), Sato et al (2007) and others. With the creation of stably labelled genetic clones in anatomically discrete areas, we hope to be able to uncover the genetic and cellular regulatory networks that govern areas of the developing embryo which have been inaccessible to labelling by other means in chicken or mouse. Here, we used our approach to comment on a long-held conundrum in limb development; the evolution of the tridactyl limb and homologies in the developing limbs of mice and chickens.

The evolution of modern birds with powered flight from basal ground-based dinosaurs has captivated the interest of scientists for more than 200 years as a premier example of a major evolutionary transformation (Brusatte, 2017). Evidence from the fossil record indicates that the path to the evolution of powered flight was likely multifactorial and piecemeal, requiring many anatomical changes in the skeletal, musculature, respiratory and integument systems (Brusatte, 2017; Brusatte et al., 2014; Dececchi and Larsson, 2009; Xu et al., 2014). Few evolutionary trends towards powered flight, however, have been as commented on or contested as the dramatic reduction from the five fingered pentadactyl hand of the basal archosaurs to the tridactyl wing of modern birds, which have drawn evidence both from palaeontological and embryological perspectives in order to understand the mechanisms by which digits were lost (reviewed Xu and Mackem, 2013). Fate-mapping to establish the origin of avian digits, as well as ascertaining the contribution of SHH signaling and *SHH* expressing cells to digits, has been used to support the evolutionary origin and therefore digit identity in modern birds, although interpretation can support both the frame-shift and axis-shift models (de Bakker et al., 2013; Kawahata et al., 2019; Tamura et al., 2011; Towers et al., 2011).

In the pentadactyl mouse limb, three main identifiers can be used to describe digit 4; articulation with the ulna as an extension of the primary axis, the first in order of appearance as a *Sox9*+ or alcian blue+ anlage over other digits (Shubin and Alberch, 1986) and derivation from *SHH* expressing cells (Harfe et al., 2004). These identifiers have also been applied to tridactyl bird wings to establish digit identity. As no digits in the chicken wing are derived from the *SHH* expressing lineage, it can be concluded that there has been a homeotic change in digit 4 identity to digit 3 via a possible change in SHH gradient (Tamura et al., 2011) or that digits 4 and 5 have been lost in the tridactyl wing evidenced by the lack of ZPA progeny found in the third digit (Towers et al., 2011), respectively. Furthermore, the tetradactyl chicken leg is concluded to have digits 1 through 4, as the fourth digit contains ZPA descendants (Towers et al., 2011).

Our finding that ZPA descendants contribute to the distal carpal 3 and posterior digit 3 metacarpal in the chicken is different from Towers et al 2011, but we believe it is due to the greater enhancement in visualisation of grafted cells through use of two transgenic reporter lines. This does not, however, change the interpretations of Towers et al (2011); digit 3 cartilage of the mouse also contains *SHH* expressing cells and by this measure we interpret that digit 3 in mouse and chicken are analogous. In summary, we find that the most posterior digit in the chicken wing is similar to digit 3 in the mouse, supporting the loss of digit 4 and 5 during evolution of the bird wing.

The zeugopod element of the primary axis, i.e. the ulna, is treated as a fixed and un-altered point, from which digit number and articulation subsequently change. Indeed, compared to digits, the ulna does superficially seem unaltered as it features in the majority of tetrapod limbs, from stem tetrapods such as the acanthostega to modern vertebrates. There are only two bones in the zeugopod; thus, its post-axial position and earlier condensation in relation to the other bone, the radius, appear to satisfy the criteria for the ulna. This principle extends to the hindlimb with the fibula recognised as analogous to the ulna (Towers, 2018). Due to the apparent conservation of the zeugopod skeleton, the distribution of ZPA descendants to the ulna of the pentadactyl limb in comparison to the tridactyl limb, or indeed to the fibula, has not yet been investigated.

In both the mouse and chicken, the ulna is dependent on SHH signaling as shown by its absence in *Shh* knockout mice (Chiang et al., 2001; Kraus et al., 2001) and OZD chicken, in which limb-specific SHH signaling is lost (Ros et al., 2003). We mapped the chick ulna in the stage HH20 limb bud, showing that it consistently arises from a highly discrete area that is adjacent to anterior somite 19 and predominantly outside of the ZPA (Fig. 5A, B). Its stage HH20 primordium is *SHH*-but *PTCH1*+, suggesting that the ulna is primarily patterned through paracrine SHH activity. Overall, the mouse and chicken ulna both appear to be largely subject to paracrine SHH signaling but have a varied distribution of *SHH* descendants.

**Figure 5.**
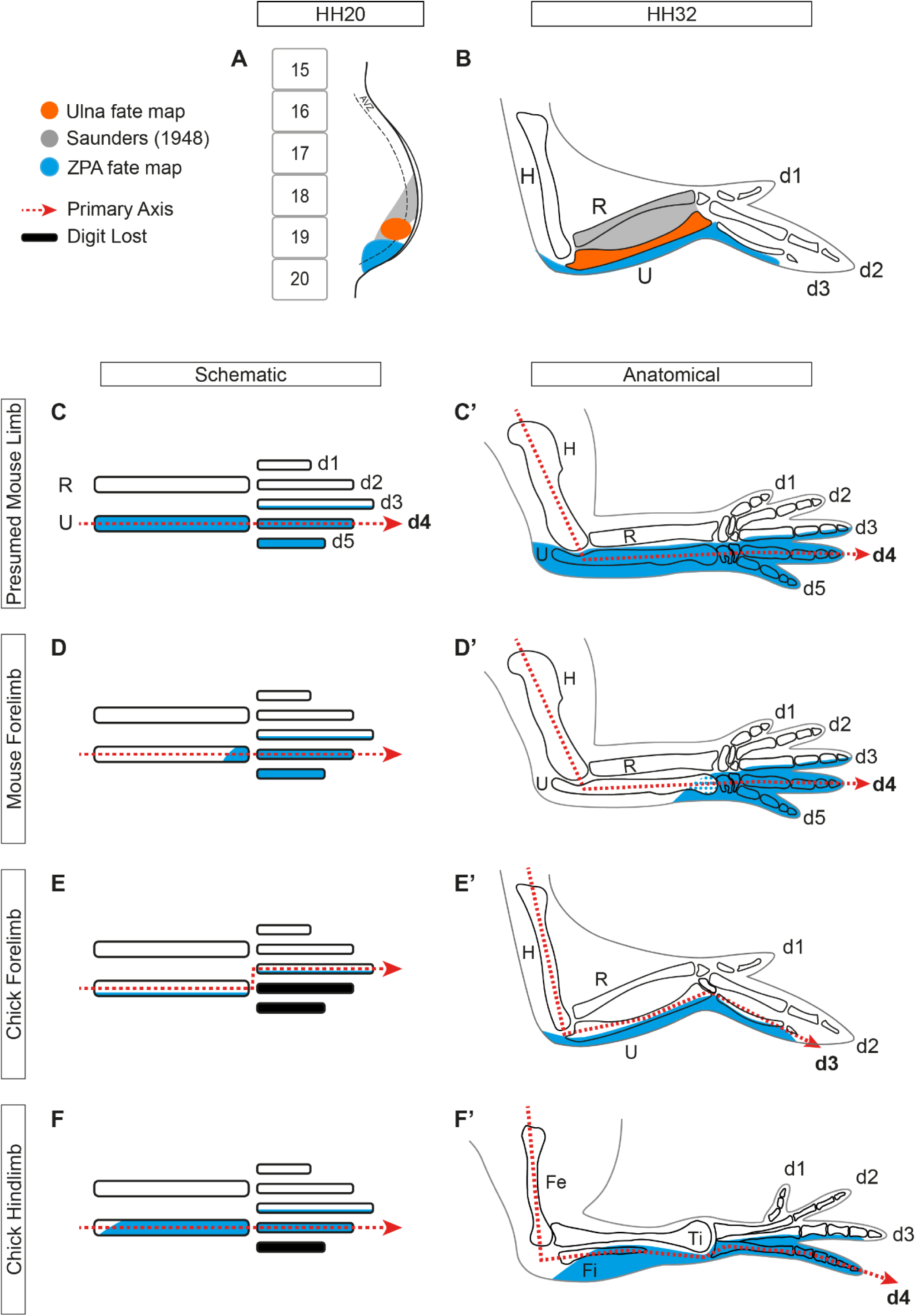
**Summary schematic of ulna fate map and ZPA lineage in relation to the primary axis** Updated fate map with the inclusion of our findings in orange (**A**, **B**). The ulna arises from the anterior half of somite 19 in the distal 20HH chick wing bud (**A**). Schematic and anatomical representations of the mouse forelimb bones with the primary axis going through the ulna and digit 4 with the presumption that the ZPA contributes to the ulna, digit 4 and digit 5 (Krawchuk et al., 2010; Zhu et al., 2022) (**C**, **C’**). Current findings that only the distal ulna of the mouse forelimb is derived from ZPA cells, demonstrating that the primary axis and ZPA lineage are not congruent (**D**, **D’**). ZPA cells contribute to the posterior-most ulna but unlike the mouse, the entire length (**E**, **E’**). If the primary axis is maintained through digit 4, then the chick wing has had no shift, maintaining the divergence of ZPA lineage to the primary axis. The fibula and digit 4 of the chick hindlimb are derived from ZPA cells (**F**, **F’**), illustrating that the ZPA lineage of the posterior zeugopod bone is not conserved, even within species. <u>Abbreviations</u> HH: Hamburger Hamilton. AVZ: Avascular zone. H: Humerus. R: Radius. U: Ulna. d1/2/3/4/5: digit 1/2/3/4/5.

In the OZD chicken, the fibula is also lost (Ros et al., 2003). However, unlike the ulna, we found that the majority of fibular cartilage is derived from SHH expressing cells, suggesting mostly autocrine SHH signaling. Digit identity as determined by ZPA contribution is often used as a fixed preliminary for proposing hypotheses for limb variations across species but also interchangeably between the fore and hind limbs. However, we show that even structures considered to be fixed and conserved like the post-axial zeugopod bone cannot be unified via the proportion of *SHH* expressing cells in its makeup.

This limitation has been acknowledged (Xu and Mackem, 2013) and through RNA sequencing of digits across five species, it has been shown that apart from digit 1, there is little homology in the expression profiles over digits 2 to 5 including that of *SHH* (Stewart et al., 2019). They conclude that the three digits of the avian wing correspond to the current amniotes expression profiles of digits 1, 3 and 4. RNA sequencing for the ulna and fibula are not yet completed but we do not expect such conservation of gene expression across species as demonstrated in digit 1 for the post-axial bone of the zeugopod. Instead, the differences in ZPA contributions between the mouse ulna, chick ulna and chick fibula indicate a composite nature of a singular anatomical structure, consisting of a complicated underlying developmental course and dynamic, piecemeal evolutionary change of its own.

Our results also suggest that the primary axis and ZPA lineage are not consistently related (Fig. 5C-F). The ulna and fibula are acknowledged to be a fixed element of the primary axis and digits have often been identified by their articulation and order of appearance in relation to the ulna or fibula (Larsson and Wagner, 2002; Shapiro et al., 2003). If there are variations in ZPA contributions between the ulna and fibula and also between species, perhaps the primary axis can also run through any digit regardless of its ZPA lineage, not just through digit 4 as has been the mainstay of digit identification. In mammals that demonstrate a reduction in SHH signaling and subsequently a loss of digits such as the pig and cow, the primary axis is maintained as with the mouse patterning, through the ulna articulating with digit 4 (Cooper et al., 2014; Tissieres et al., 2020). However, in birds with a delay and reduction in relative SHH signaling, which cause a loss of posterior digits and carpals such as the emu, the ulna articulation shifts anteriorly to digit 3 (Kawahata et al., 2019; Smith et al., 2016), suggesting digit identity as determined by SHH lineage does not dictate the course of the primary axis.

In conclusion, we show although the mouse and chicken ulna are predominantly *SHH*-, suggesting paracrine patterning. Unlike the ulna, chick fibular cartilage is mostly descended from the ZPA and thus, although the postaxial zeugopod is seen as fixed and often considered as analogous, we demonstrate that these actually have different constituents of SHH lineage. The ulna and fibula may be more evolutionarily diverse than supposed and therefore, their participation in the primary axis may be flexible and unrelated to ZPA lineage. We suggest that with changes in digit number, the articulation of the zeugopod with the autopod have correspondingly developed to accommodate functionality over digit identity and that the zeugopod will have adapted just as much as digits, alluded to by the variation in contributions of *SHH* expressing cells.

## Materials and Methods

### Chicken Husbandry

ISA Brown, Roslin Green (Cytoplasmic GFP), Flamingo (TdTomato) and Chameleon (Cytbow) chicken lines were maintained under Home Office License at the Roslin Institute. Fertilised chicken eggs were incubated at 38°C until the desired stage of embryonic development (Hamburger and Hamilton, 1951).

### Mouse Construction and Genotyping

Mice used in this study were housed at the animal facilities at the University of Edinburgh, with procedures performed under Personal and Project Home Office Licences. Male mice carrying the SHHtm1(EGFP/cre)Cjt allele (Harfe et al., 2004) were mated to female mice carrying a Cre reporter line (Luche et al., 2007). Cre expression leads to excision of a floxed transcriptional Stop cassette and allows expression of the tdRFP in all descendant cells. Embryos were collected at E14.5, genotyped by standard methods and fixed overnight in 4% PFA.

### Homotopic Grafts

At the desired stage, host sites of Roslin Green or ISA Brown embryos were dissected and discarded using a tungsten dissecting needle. Donor sites from Flamingo embryos were dissected and moved into the host Roslin Green embryo via a p20 pipette containing DMEM. The graft was manoeuvred into the host site and, when necessary, secured with a piece of 0.02mm oxidised nickel chrome wire. Care was taken to ensure ectoderm orientation was maintained between donor and host. Embryos for wholemount analysis were culled and dissected at around stage 33HH, fixed and cleared with CUBIC reagent 1 before being imaged on a Zeiss Axiozoom V16 microscope. Embryos for HCR in situ hybridisation were allowed to incubate for 3 hours after graft insertion, then culled and dissected in cold DEPC PBS before being fixed with 4% PFA at 4°C overnight.

### Chameleon Cytbow Chicken Manipulations

Fertilised eggs were windowed, prepared for manipulation as per Tiecke and Tickle (2007) and staged (Hamburger and Hamilton, 1951). Once bead manipulations were complete, the window was sealed with tape and incubated at 38°C in a humidified and light-free environment until the desired Hamburger and Hamilton stage. Embryos were culled in accordance with Schedule 1 of the Animals (Scientific Procedures) Act 1986. Embryos were dissected in cold PBS in preparation for staining or in-situ hybridisation.

### Hybridisation Chain Reaction In Situ Hybridisation

Whole-mount tissue was prepared for HCR by dissecting in cold DEPC PBS and fixing in 4% PFA overnight. After washing twice in PBT for 5min each, fixed tissue were dehydrated with a series of MeOH/PBST washes for 5min each on ice. Once dehydrated up to 100% MeOH, tissue were stored in -20°C until further use. Prior to performing HCR, tissues were rehydrated with a series of MeOH/PBST washes for 5min on ice up to 100% PBST. Tissues were treated with 10ug/mL proteinase K solution at room temperature for a length of time that was calculated at 15sec per stage (e.g. 5min for stage 20HH). These were post-fixed in 4% PFA at room temperature, then washed twice in PBST for 5min each, 50% PBST/50% 5XSSCT for 5min, then 5XSSCT for 5min, all on ice. We then performed HCR v3.0 using the protocol as described by Molecular Instruments (Choi et al., 2018). Split initiator probes (v3.0) for *PTCH1* (accession #NM_204960.2) and *SHH* (accession #NM_204821.1) were designed by Molecular Instruments, Inc.

### Sections

Embryos were dissected in cold PBS and fixed in 4% PFA overnight. After sucrose treatment, limbs were embedded in a solution of 7.5% gelatin and 15% sucrose in PBS then frozen in isopentane at around -60°C and stored at -80°C until sectioning. Serial sections were obtained with a Bright OTF5000 cryostat microtome at a 10um thickness and mounted on Polysine Adhesion microscope slides. Once dry, slides were washed in PBS at 37°C and mounted with coverslips. Images were obtained on the LSM880 Confocal microscope using Zen Black software.

### Clearing

Fixed tissues were washed in PBS for 5min at room temperature then submerged in CUBIC reagent 1A (as per Susaki et al., 2015) at 37°C for 2-6 hours until cleared.

### PCR

Five samples of ulna and ZPA were dissected from 20HH ISA Brown embryos and batched for a single reaction. These were stored at -80°C before RNA extraction using Pre cellys bead homogenisation (Bertin Technologies, France) and RNA easy Kit (Qiagen). Turbo DNA free DNase kit (Ambion) was used to remove genomic DNA contamination before cDNA was synthesised using AffinityScript Multiple Temperature cDNA Synthesis Kit (Agilent) using Oligo DT. Triplicate qRT-PCR reactions were carried out per biological replicate using an MX 3005P thermal cycler (Agilent) using a FAST 2 step thermal cycling protocol (95°C 10 sec, 60°C 30 sec). Brilliant iii Ultra Fast SYBR green qPCR master mix (Agilent) and Chicken primers were used at 100nM final and were as follows: LBR F: GAAGCTGCAGTACCGGATCA, LBR R: GCTAGGTCTTCCTCAGGTGC (housekeeping gene). SHH (accession #NM_204821.1) F: CCAAATTACAACCCTGAC, SHH R:CATTCAGCTTGTCCTTGCAG, PTCHD1 F: TGGGAAATACAATTCCACCTTC, PTCHD1 R: CTCCAGGAGGACAACATTTCA. Data was analysed using MX Pro software and exporting to Excel where a 2-ddCT method was used to calculate relative expression compared to ZPA.

### Contributions

Conceptualization: MGD and JDHO

Methodology and resources: MGD, JDHO, DDZS, LM, MJ, JG, JJS, LAL

Analysis and investigation: MGD, JDHO, MJ, JG

Writing – Original Draft: MGD and JDHO/All authors contributed to manuscript review and editing.

## Acknowledgments

We thank the National Avian Research Facility, The Roslin Institute and R(D)SVS for the maintenance and production of fertile eggs from the Chameleon, dtTomato, GFP lines. This study was funded by Biotechnology and Biological Sciences Research Council (BBSRC) to The Roslin Institute, BB/X010937/1, BB/CCG2270/1 and to the Institute of Genetics and Cancer by the Medical Research Council (MRC) University Unit programmes MC_UU_00007/8 and MC_UU_00035/7. JDHO is supported by a University of Edinburgh Principals Studentship and The Roslin Institute.

## Ethical Considerations

All animal experiments were reviewed and approved by the University of Edinburgh Animal Welfare and Ethics Committee and were conducted with appropriate licensing under Animals (Scientific Procedures) Act 1986. All experiments on chicken embryos were undertaken for day 14 of incubation.

