## Supplementary Figures for "Insights into Digit Evolution from a Fate Map Study of the Forearm"

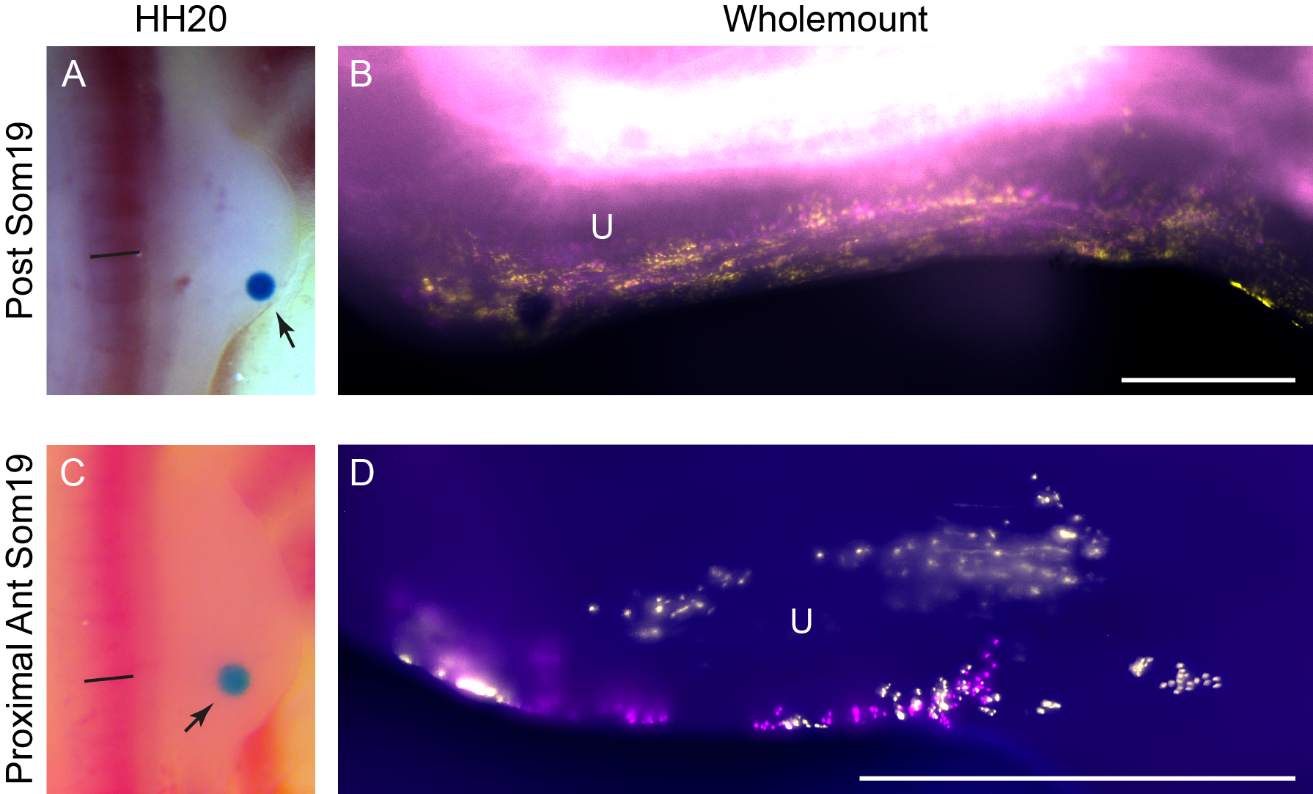

**Supplementary Figure 1. The ulna arises from a highly localised area of the 20HH chick wing bud**

TAT-Cre soaked beads (arrow) placed adjacent to posterior somite 19 in 20HH CPX chick wings (**A**) will not give rise to the ulna (**B**). Whilst adjacent to anterior somite 19, beads (arrow) placed more proximally (**C**) will also not give rise to the ulna (**D**). Straight black lines denote the anterior-most edge of somite 19. Abbreviations HH: Hamburger Hamilton. U: Ulna. All scale bars = 200μm

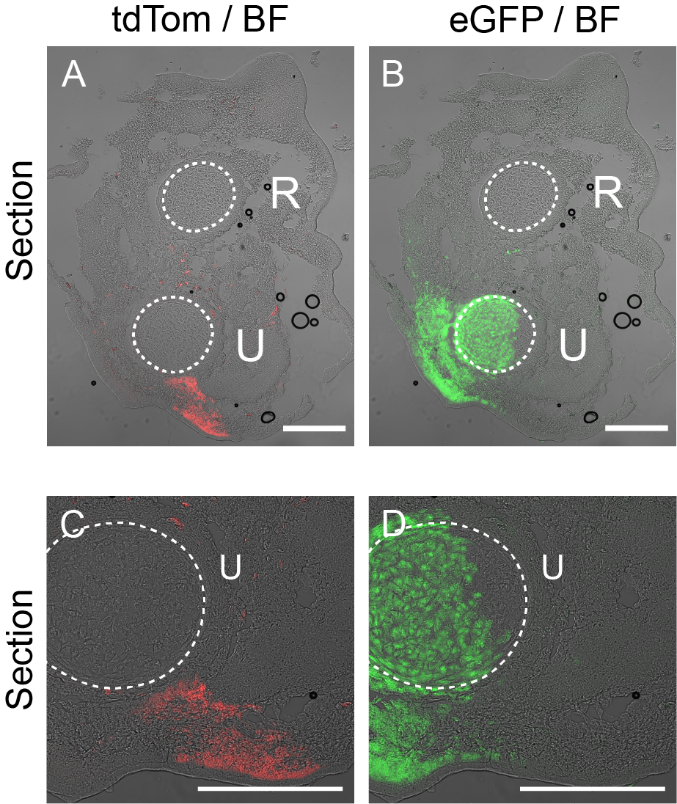

**Supplementary Figure 2. ZPA derived cells do not contribute to the ulna.**

Split channels from Fig. 3K to show no contribution of ZPA derived tdTom cells in the ulna.
